# Identification and characterization of potent and selective inhibitors for the B^0^AT2/SLC6A15 amino acid transporter

**DOI:** 10.1101/2025.03.25.645215

**Authors:** S Cuboni, F Hausch

## Abstract

The gene SLC6A15 encodes the protein B^0^AT2, a transporter for neutral amino acids. It is highly expressed in the brain and has been associated with depression but little is otherwise know about its function. In this study, we identified the first inhibitors of this protein to pharmacologically investigate its function.

The transporter activity was evaluated using a cellular uptake of the substrate ^3^H-proline. A miniaturized assay was developed and used for a High Throughput Screening (HTS) of 200,000 compounds. Hits were tested for cell toxicity and selectivity *versus* related transporters of the SLC6 family. The most promising inhibitors were validated by proline uptake and neurite outgrowth in primary hippocampal neurons.

Of the 10 chemical scaffolds identified, a 1,5-benzodiazepine series had the most promising selectivity and structure-activity relationship (SAR) profile. The best compounds showed drug-like properties and inhibited B^0^AT2 with an IC_50_ of 250 nM both in SLC6A15-overexpressing HEK293 cells and primary neurons, with no detectable inhibition (> 80µM) of SERT, DAT, GAT1, or NTT4/SLC6A17. These compounds also dose-dependently stimulated neurite outgrowth in primary neurons.

The identified compounds are the first inhibitors of the amino acids transporter B^0^AT2/SLC6A15. Their potency, selectivity and physicochemical properties allow to target the transporter in relevant biological systems and to initiate new studies to understand its role and implication in diseases.

## 2. Introduction

The SLC6 family represents one of the most important classes of transporters (Bröer *et al*., 2012; Chen *et al*., 2004), which play an important role in many physiological mechanisms and especially for psychiatric disorders (Hahn and Blakely, 2007). The SLC6 family contains the transporters of the neurotransmitters serotonin (SERT), norepinephrine (NET), dopamine (DAT), GABA (GAT1) and glycine (GlyT1), which are the targets of antidepressants, anti-convulsants, illicit drugs and of potential antipsychotics (Broer, 2006).

B^0^AT2 (also known as v7-3, NTT73 or SBAT1, endoced by the SLC6A15 gene) is a new member of the SLC6 family, which is highly expressed in different brain regions including the olfactory bulb, cerebral cortex, cerebellum hypothalamus and hippocampus (Farmer *et al*., 2000; Hägglund *et al*., 2013; Inoue *et al*., 1996). The SLC6A15 mRNA and protein are also highly expressed in neurons and astrocytes in the hypothalamus (Hägglund *et al*., 2013). Like other members of the SLC6 family B^0^AT2, the protein encoded by SLC6A15, is a Na^+^-coupled symporter. It mediates the transport of neutral amino such as proline, methionine, isoleucine and other branched-chain amino acids (Broer *et al*., 2006; Drgonova *et al*., 2007; Takanaga *et al*., 2005).

Recently, a genome-wide association study (GWAS) identified SLC6A15 as a possible susceptibility gene for Major Depressive Disorder (MDD) (Kohli *et al*., 2011). The authors showed a genetic correlation between a single nucleotide polymorphism (SNP) rs1545843 downstream of the human SLC6A15 (hSLC6A15) SNPs and increased stress susceptibility in depressive patients. Patients with MDD, which carry the risk allele, also have higher adrenocorticotropic hormone (ACTH), higher cortisol responses and a lower cognitive performance suggesting a role of the SLC6A15 transporter on hypothalamic-pituitary-adrenocortical (HPA) axis regulation (Schuhmacher *et al*., 2013). Moreover, another genetic study identified two rare non-synonymous variants of the SLC6A15 transporter, which were associated with an increase of maximal proline uptake compared to the wildtype sequence (Quast *et al*., 2013).

Altogether, these results point to a putative involvement of the B^0^AT protein in psychiatric disorders. In order to better understand the function of B^0^AT2 we developed a HTS assay and used it for the identification of the first inhibitors for this transporter.

## 3. Methods

### 3.1 Materials

The HTS was performed using the COMAS screening library with holds ca. 200,000 compounds including 140,000 commercial compounds (ChemDiv, Enamine), natural-product derived compounds of the Waldmann collection (Max Planck Institute of Molecular Physiology, Dortmund) and the compound collection of the Lead Discovery Center (45,000 compounds). Compounds for hit confirmation and of the analogs of family 10 where obtained from Ambinter or from AKoS GmbH and had > 95% purity.

#### Cloning and stable clone selection

The cDNA of the long form of SLC6A15 (IRAKp961L15168Q; ImaGenes) was inserted into the pEGFP-C1 vector (Clontech) using the restriction sites BgIII and SalI. A stable SLC6A15-overexpressing HEK293 cell line was generated using the selection marker geneticin.

HEK293 cells stably expressing the transporters hSERT and hDAT were kindly provided by Blakely et al (Qian *et al*., 1997) and the HEK293-hGAT1 cell line was provided by K. Wanner (Kragler *et al*., 2008). The plasmid NTT4-B^0^AT2 (the carboxyl terminus of NTT4 was replaced with the analogous region of B^0^AT2) was kindly provided by the Reimer lab (Zaia *et al*., 2009). This plasmid was used to generate a cell line HEK293 stably expressing the NTT4-B^0^AT2 chimera.

### 3.2. Cell culture and cryo-cells

HEK293 eGFP-hSLC6A15 cells were cultured in Dulbecco’s Modified Eagle medium (DMEM, Gibco) containing 10 % of fetal calf serum (FCS), 500 units/ml penicillin and 500 µg/ml streptomycin at 37ºC in a humidified incubator (5 % CO_2_). For the screening cryo-cells were prepared. HEK293 eGFP-hSLC6A15 cells were plated in 10-stack CellSTACK chambers (Corning). After harvesting, aliquots of 2 x 10^7^ cells/ml in complete medium containing 10 % DMSO were stored in liquid nitrogen until use.

### 3.3. Preparation primary neurons and virus transfection

Hippocampal neurons were prepared from wild type C57/BL6 mice at embryonic day 18. Hippocampi were removed from the whole brain and transferred to the dissection medium (HBSS containing Ca^2+^, Mg^2+^ and phenol supplemented with 50µM *D*-2-amino-5-phosphonopentanoate (D-AP5)). Cells were dissociated with different pipetting steps and centrifuged for 5 minutes at 200 g. The pellet was resuspended in plating medium (DMEM with L-glutamine, phenol red, pyruvate, without HEPES and supplemented with 5 % FCS and 1 % GlutaMAX-I (Gibco)) and plated. After 4 hours, medium was replaced with growing medium (Neurobasal medium supplemented with 2% B27, 1 % GlutaMAX-I, 50 units/ml penicillin and 50 µg/ml streptomycin). Cells were incubated at 37 ºC in a humidified incubator (5 % CO_2_) for 11 days. 3 days before the uptake assay cells were transfected with 0.4 x 10^9^ genomic particles/ml of the adeno-associated virus AAV1/2-CAG-Mouse SLC6A15-WPRE-BGH-polyA or of the corresponding empty vector AAV1/2-CAG-Null/Empty-WPRE-BGH-polyA (GeneDetect^®^).

### 3.4. _3_H-proline uptake assay in primary neurons and HEK eGFP-hSLC6A15

For assay development and for characterization in primary neurons the ^3^H-proline uptake assay was performed in 96-well plate (Corning 3610) using 60,000 cells per well. For screening and profiling 384-well plates (Corning 3846) with 10,000 cells per well were used. HEK eGFP-hSLC6A15 cells were dispensed in plates pre-coated with poly-D-lysine (PDL) and incubated overnight at 37 ºC in a humidified incubator (5 % CO_2_). After 24 hours, the medium was removed with six washing steps (ELx405, Biotek) and replaced with the assay buffer (4.7 mM KCl, 1.2 mM MgSO_4_, 2.2 mM CaCl_2_, 0.4 mM KH_2_PO_4_, 10 mM D(+)Glucose, 10 mM HEPES, 120 mM NaCl, pH 7.4) (Zaia *et al*., 2009). Compounds dissolved in DMSO were added (Echo 550, Labcyte) and incubated at room temperature (RT) for 10 minutes. The final DMSO concentration in the assay was 0.8 %. Afterwards, 125 nM ^3^H-proline was dispensed (Microdrop, Thermo Scientific) and cells were incubated for other 10 minutes at RT. The uptake reaction was stopped by nine washing steps performed with assay buffer. Cells were lysed with 1 M NaOH and the radioactive signal amplified with scintillation cocktail (PerkinElmer). Plates were incubated for 6 hours in the dark and the radioactivity was quantified using a Wallac Microbeta (PerkinElmer) by counting for 2 minutes per well.

The uptake assays in HEK-NTT4/B^0^AT2, HEK-hSERT, HEK-hGAT1 and HEK-hDAT cells were performed in 384 well plates following the protocol for HEK-eGFP-hSLC6A15 using transporter-specific buffers. For ^3^H-5HT uptake in HEK-hSERT the assay buffer was composed of 5 mM KCl, 1.2 mM MgSO_4_, 2.5 mM CaCl_2_, 11 mM D(+)Glucose, 25 mM HEPES, 120 mM NaCl, pH 7.4. Before usage 1 µM of pargyline was added (Deecher et al., 2006). For ^3^H-GABA uptake in HEK-hGAT1 the assay buffer used was the same as for the proline uptake with 25 mM Tris. For ^3^H-dopamine uptake in HEK-hDAT the assay buffer was composed of 4.7 mM KCl, 1.2 mM MgSO_4_, 2.2 mM CaCl_2_, 1.2 mM KH_2_PO_4_, 10 mM HEPES, 120 mM NaCl, pH 7.4 (Riherd et al., 2008). Paroxetine, tiagabine, dopamine, and cold proline were used as positive controls in the SLC6A transporter specificity screens, respectively.

### 3.5. Measurement of neurite outgrowth

Dissociated primary hippocampal neurons from 18 days old C57/BL6 mouse embryos were plated in 12-well plates containing precoated coverslips at a densitiy of 100,000 cells/well. After 4 hours, the plating medium was replaced with growing medium containing the compounds and cells transfected with an adeno-associated virus coding for GFP (GeneDetect^®^). The assay was performed using 0.005 % DMSO as final concentration. After 48 hours, the medium was removed and cells were fixed with 4 % paraformaldeyde (PFA) for 20 minutes. For better visualization of the GFP, cells were permeabilized three times for 5 minutes with Triton-X 100 0.1 % and than immunostained overnight at 4 ºC with an anti-GFP antibody (Abcam). The antibody was than washed three times with PBS and cells were incubated with an AlexaFluor 488-conjugated secondary antibody (Invitrogen). After 2 hours of incubation time at room temperature, cells were washed and coverslips transferred to object slides for imaging.

### 3.6. Microscopy and neurite outgrowth analysis

Imaging of the cells was performed using a fluorescence microscope (Axiolan 2) with 40 fold magnification. At least 30 cells per condition were analysed and the total neurite length was determined using ImageJ software with NeuronJ plugin (NIH).

### 3.7. Cell viability assay

1000 cells were plated in low volume plates (Corning 3570) in a total volume of 12.5µl per well. After 24 hours, cells were treated with the compounds (0.7 % DMSO final concentration) and incubated for after 24 hours. The cell viability assay was performed following the protocol provided by Promega for CellTiter-Glo^®^ Luminescent Cell Viability assay.

### 3.8. Statistical Analyses

Data are given as mean S.E.M. For statistical comparison using PRISM_4_, Students t*-*test was used. p values less than 0.05 were considered statistically significant. (*p < 0.05, **p < 0.01, ***p < 0.001).

## 4. Results

### 4.1. Assay development

HEK293 cells stably over-expressing eGFP-hSLC6A15 were generated and the correct plasma membrane localization of B^0^AT2 was confirmed (Fig. 1a). To assess the amino acid transport activity a ^3^H-proline uptake assay was established in a 96-well plate format and key experimental conditions were optimized. ^3^H-proline uptake was saturable with a K_M_ value of 960 ± 51 µM and a V_max_ of 3838 ± 66 pmol/min/10^6^ cells (Fig. 1b). The uptake increased linearly in the first 20 minutes and then approached a plateau after 40 minutes (Fig. 1c). As described, B^0^AT2 transporter activity was strictly sodium-dependent and strongly compromised under acidic conditions (Fig.1d) (Broer *et al*., 2006; Takanaga *et al*., 2005). B^0^AT2 activity was not affected by DMSO concentrations up to 2 %. A pH = 7.4, a DMSO concentration of 0.8 %, 80 nM ^3^H-proline and a reaction time of 10 minutes were chosen for screening. Using these conditions, inhibition curves were performed with unlabelled proline and leucine, which inhibited B^0^AT2 with an IC_50_ of 257 ± 53 µM and 210 ± 30 µM, respectively (Fig. 2). In order to identify small molecules that could modulate the B^0^AT2 activity in a high-throughput manner, the optimized conditions were adapted to a 384-well plate format using a semi-automated workflow. A Z’-factor of 0.78 was determined showing the high robustness of the assay (Zhang *et al*., 1999).

**Fig. 1:**
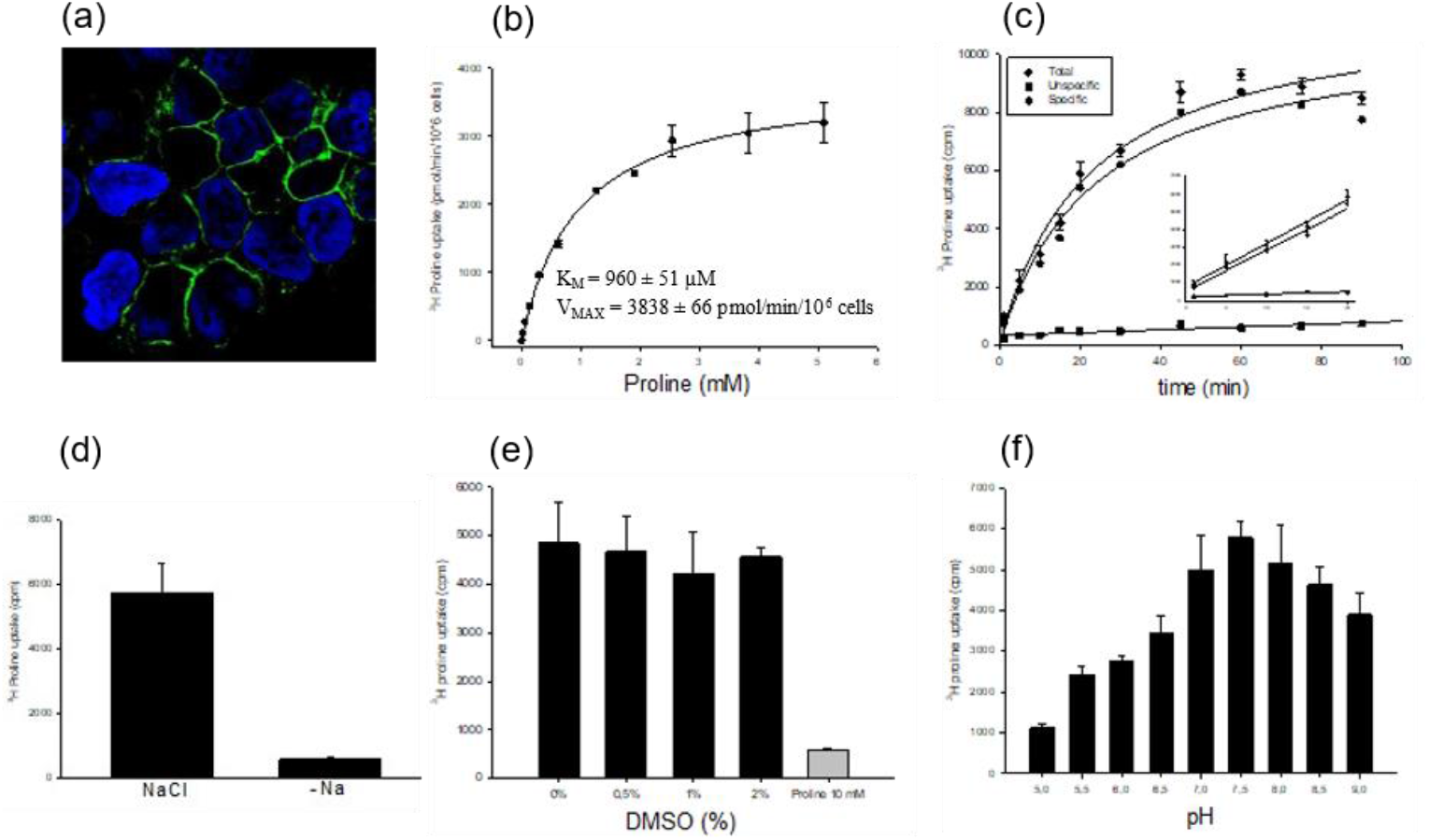
Assay development for the screening of B^0^AT2/SLC6A15. (a) HEK293 cells stably expressing hSLC6A15 as an eGFP fusion showed the expected plasma membrane localization. (b) The saturation analysis of B^0^AT2/SLC6A15 was performed by HEK293 eGFP-hSLC6A15 cells with increasing concentrations of ^3^H-proline. (c) The ^3^H-proline uptake increased linearly in the first 20 minutes and than approached a plateau after 40 minutes. Unspecific uptake was determined in the presence of 10 mM unlabelled proline. (d) B^0^AT2 is a sodium coupled transporter and removal of sodium (120 mM) from the assay buffer led to a complete block of the amino acid uptake. (e) The uptake was not significantly affected by DMSO concentrations up to 2 %. (f) pH dependence of B^0^AT2. ^3^H-proline uptake was optimal at pH = 7.5.

**Fig. 2:**
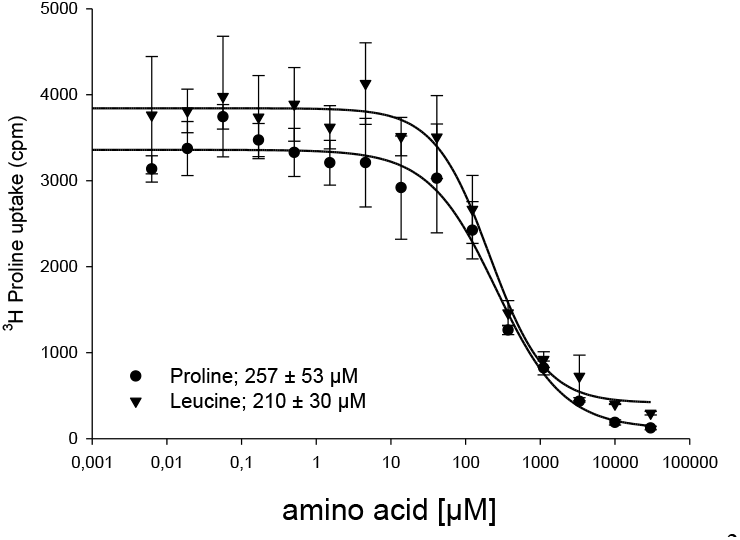
Proline and leucine inhibition curves. ^3^H-proline uptake of hSLC6A15 was performed in presence of increasing concentrations of unlabelled proline or leucine.

### 4.2. Screening and Hit Validation

The ^3^H-proline uptake assay was used to screen a library of 200,000 diverse substances at a final compound concentration of 10 µM. A cut-off of 30 % residual activity was selected yielding 126 validated primary hits (hit rate = 0.06 %). These were retested in dose-responses curves and counter-screened for cell toxicity. The remaining confirmed hits were clustered in 10 scaffolds based on the chemical structure (Tab. 1).

**Tab. 1.**
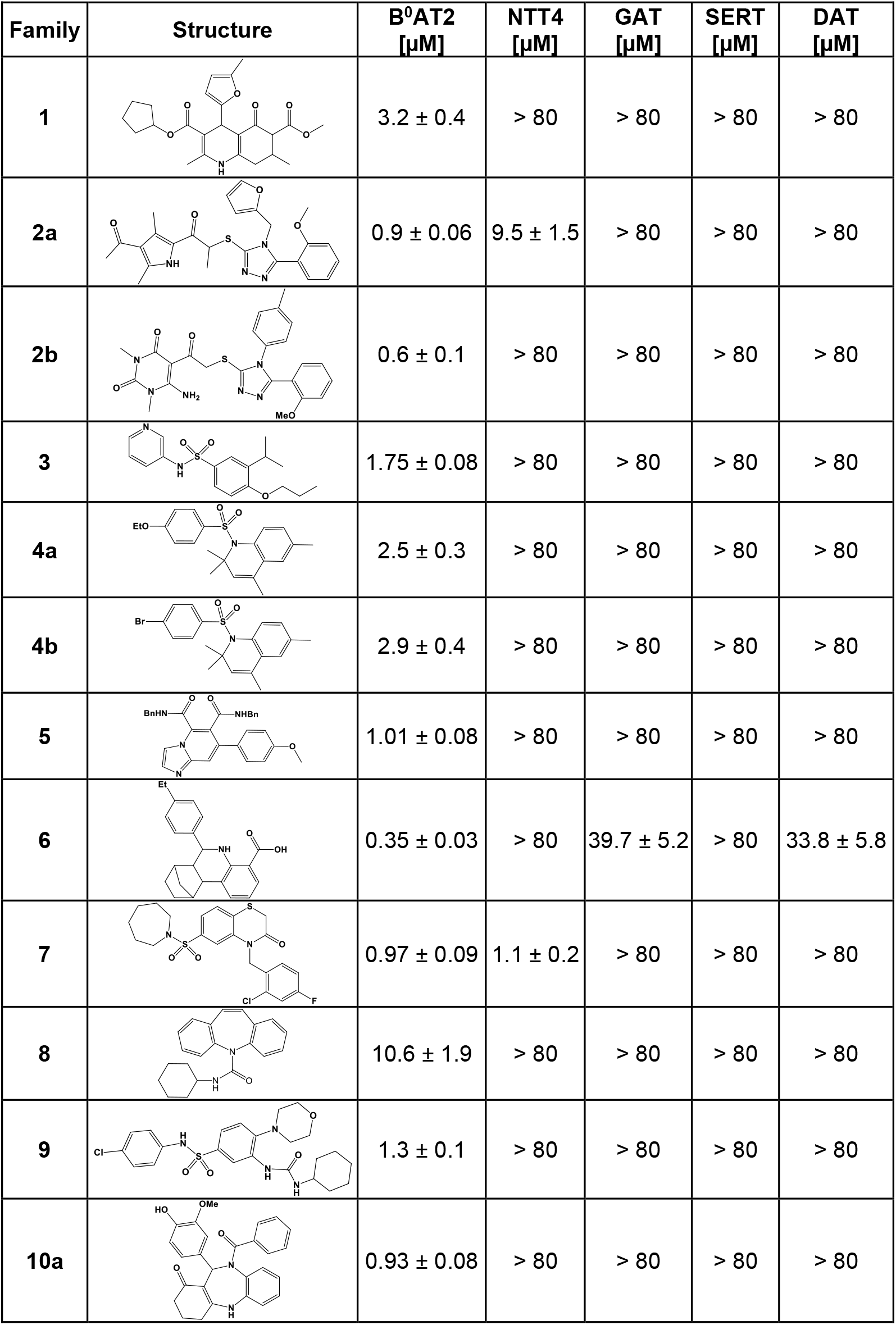
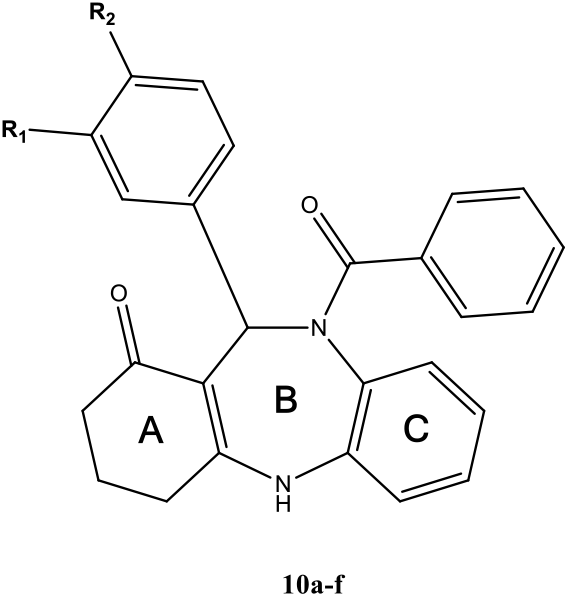

The confirmed hits were profiled for selectivity against the transporters for GABA (hGAT1), serotonin (hSERT), dopamine (hDAT) and NTT4/SLC6A17, the closest homolog of B^0^AT2/SLC6A15. The results pointed to the 1,5-benzodiazepine **10a** as being the most promising B^0^AT2-selective scaffold, which was explored in more detail.

### 4.3. Structure activity relationship (SAR) of the 1,5-benzodiazepine scaffold

We first probed the substitution pattern at the exocyclic phenyl ring of hit **10**. This indicated that diverse small groups where tolerated in the meta and para position (Tab. 2). This was confirmed in a more extensive SAR analysis of the 1,5-benzodiazepine scaffold **11**, which had an additional geminal dimethyl group in the A-ring. Both electron-donating and electron-withdrawing substituents were tolerated in the para-position (**11a-m**, Tab. 3), while larger residues like t-butyl (**11n)** were inactive. In the meta-position, fluorine reduced activity while ether and ester groups were tolerated (**11o-r**). Any substituent in the ortho-position, however, substantially reduced activity (**11s-u**). Meta-/para-bi- or tri-substitution did not enhance activity (**11w-ad**), with the exception of **11v**, which was one of the most potent B^0^AT2 inhibitors. Both 5- and 6-membered heterocycles could also replace the exocyclic aromatic group of the B-ring, albeit usually with reduced potency (**12a-i**, Tab. 4). In contrast, the benzoyl substituent of the B-ring was absolutely critical. Any modification other than a fluorine substitution abolished inhibition of B^0^AT2 (Tab. 5). Finally, we explored substituents in the 3-position of the A-ring (Tab. 6). Para-substituted phenyl groups and a furanyl heterocycle where possible (**14a**,**b**,**d**,**e**,**g**), whereas a meta-/para-di-substituted phenyl group was not (**14f**). However, in the context of a different substitution pattern at the exocyclic phenyl group of the B-ring several C-ring substituted analogs were inactive (**14h-l**). This suggests that the exocyclic aromatic moieties of the B- and C-rings interact with each other in the binding site of B^0^AT2. All 1,5-benzodiazepine analogs were tested for inhibition of SERT, GAT1, SLC6A17 and for cell toxicity. No activity for any of these transporters (> 80 µM) and no sign of cell toxicity could be detected for any of these compounds. Several representative 1,5-benzodiazepine analogs including compounds **11x, 14c** and **14d** were also tested for inhibition of DAT, which were all inactive for this transporter (> 80 µM).

**Tab. 2.**
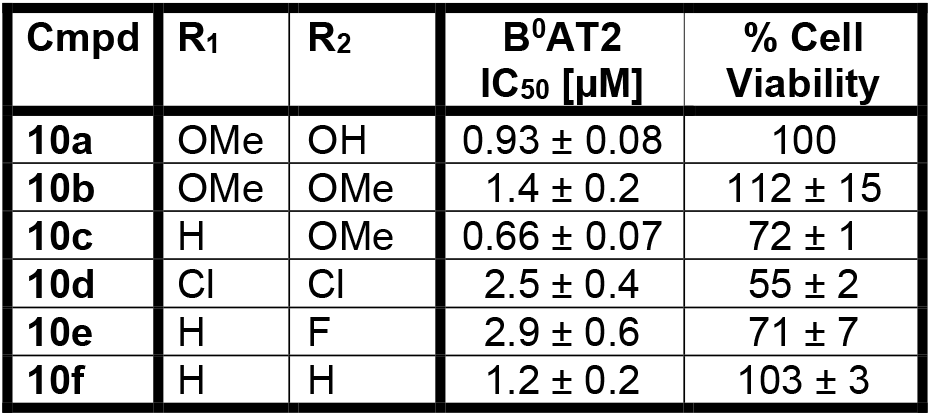
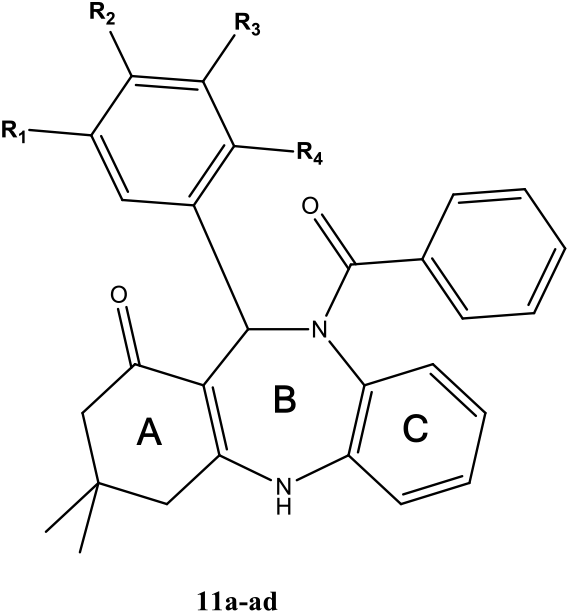

**Tab. 3.**
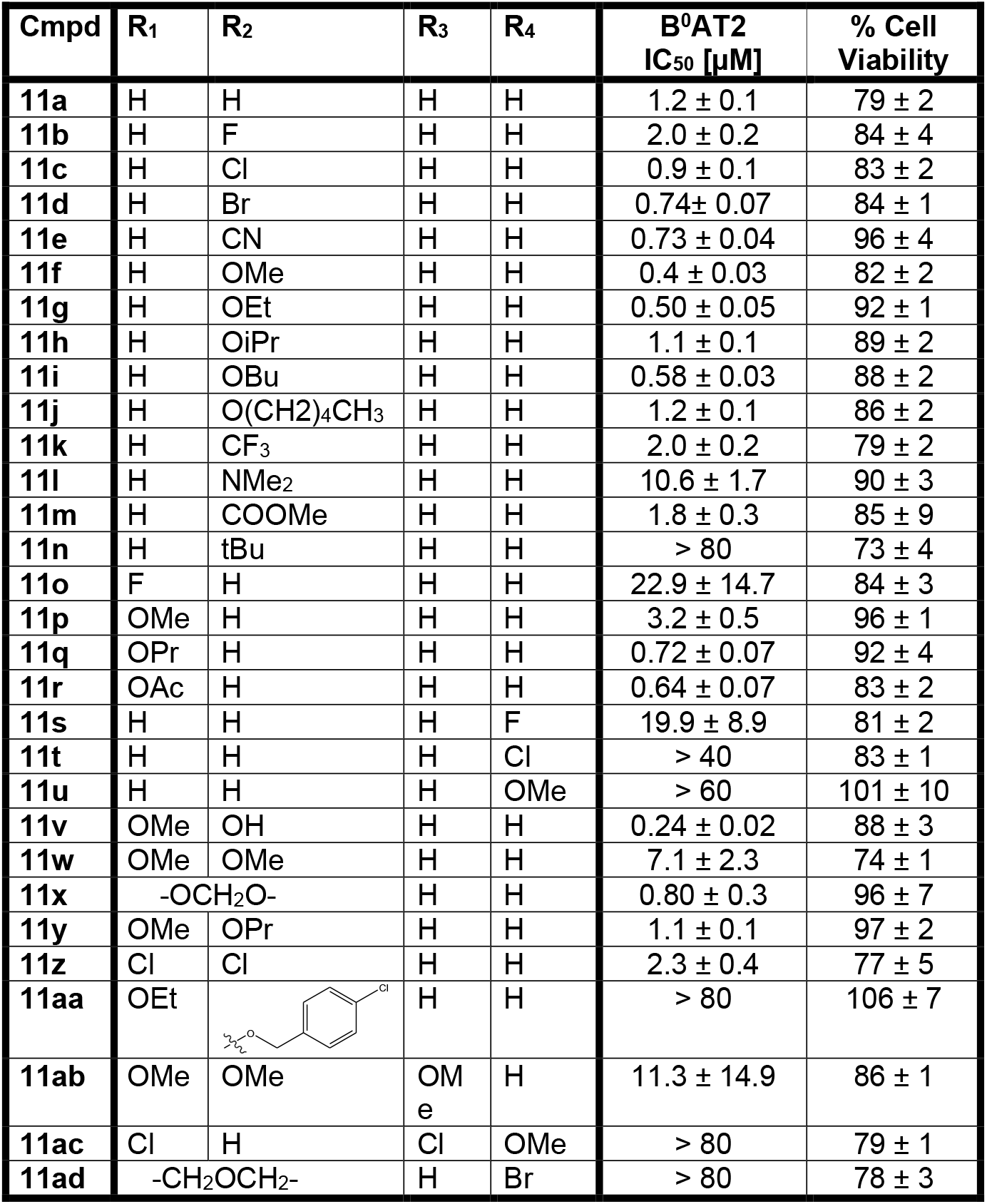
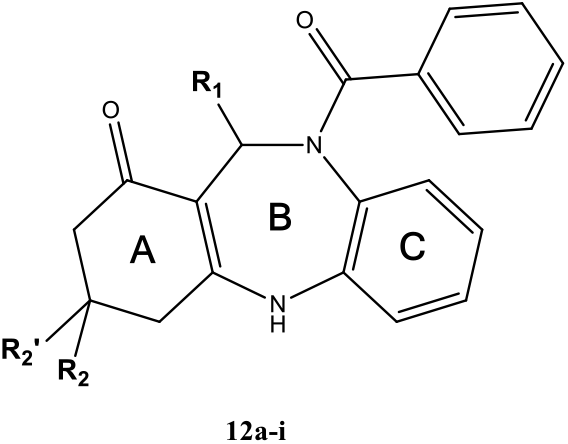

**Tab. 4.**
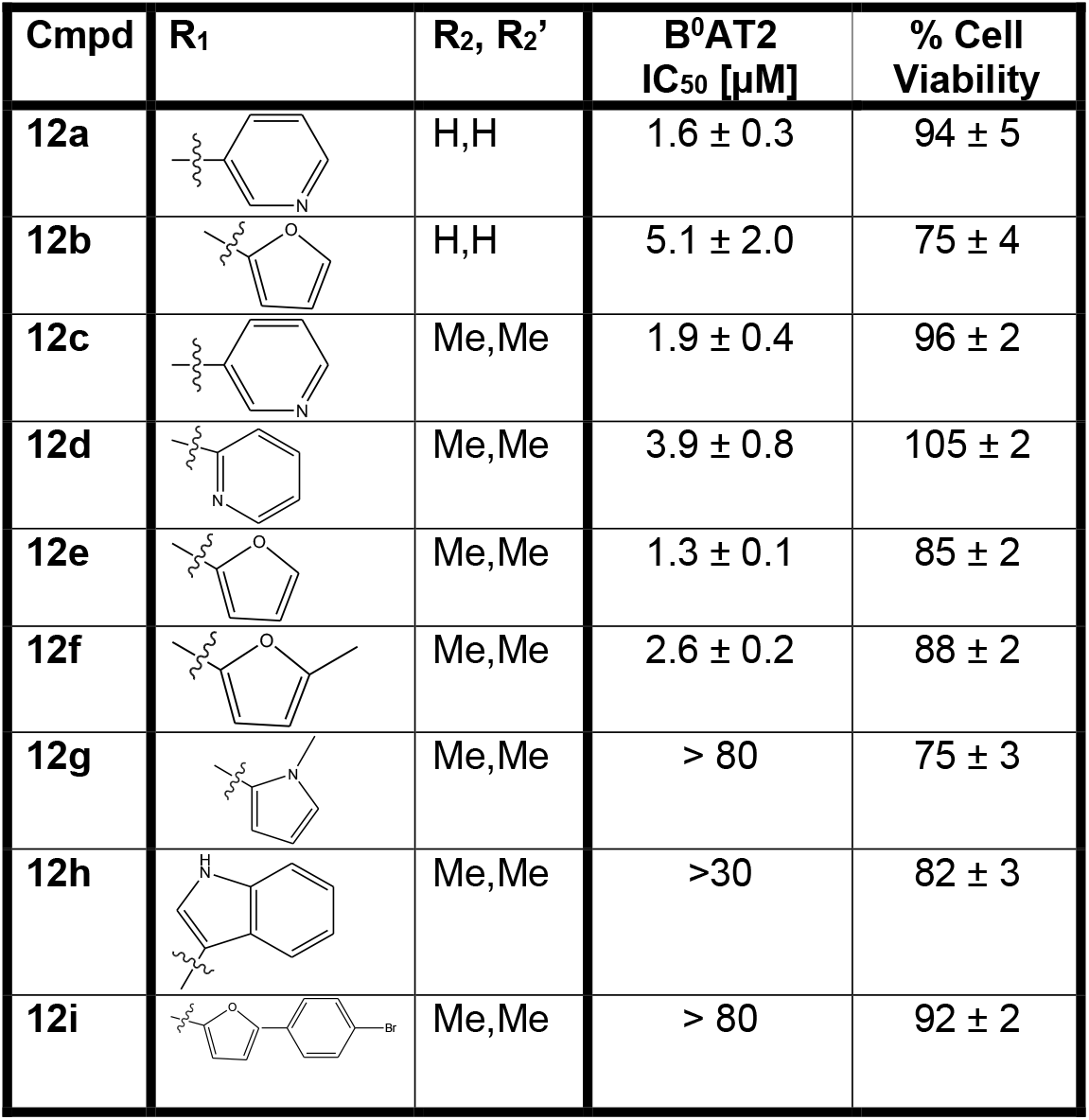
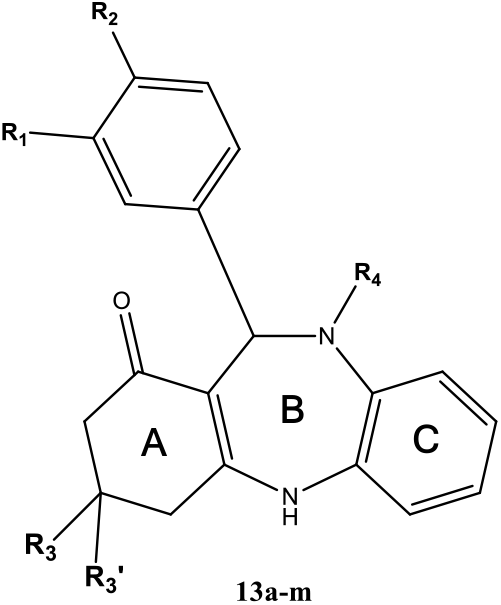

**Tab. 5.**
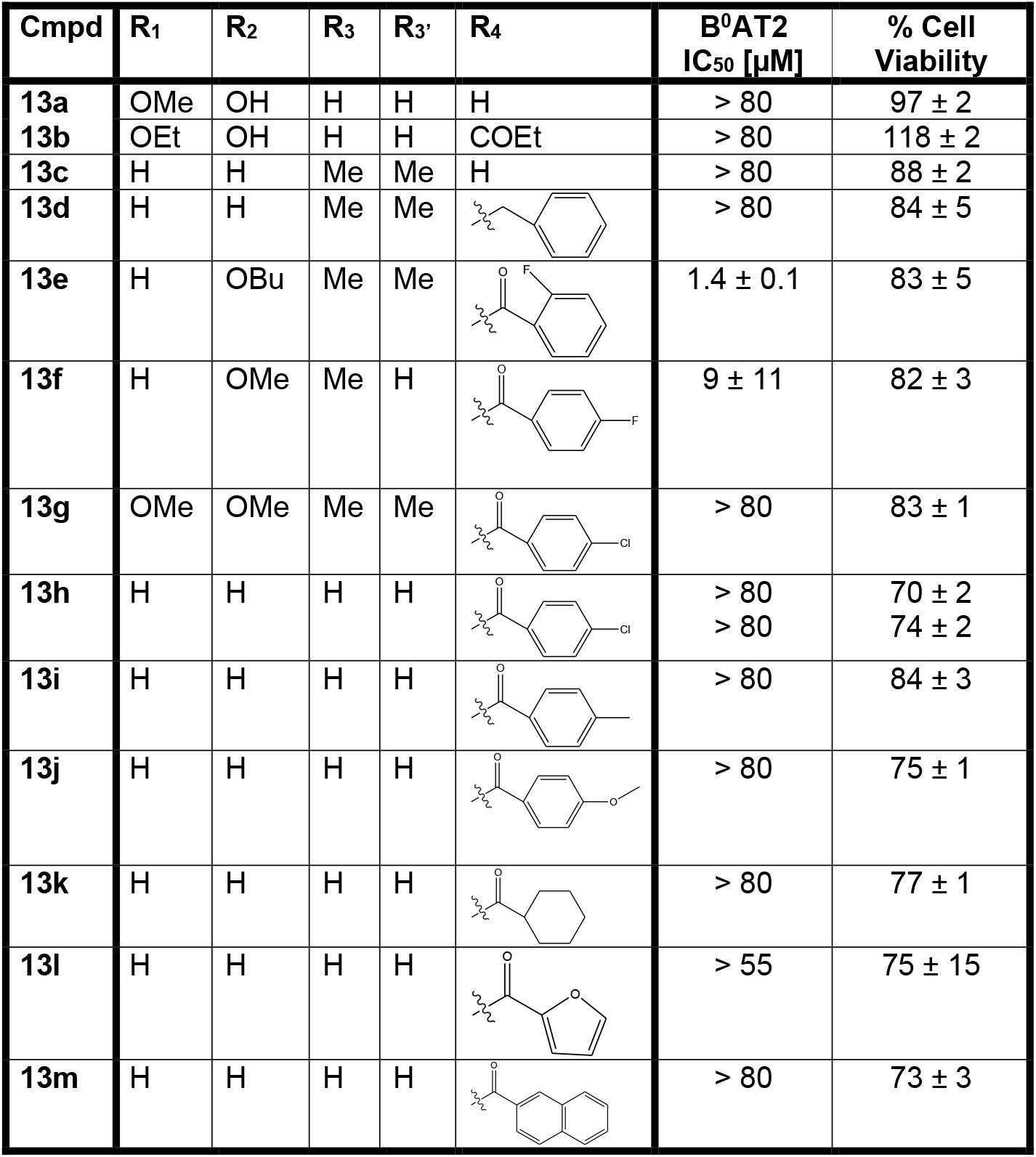
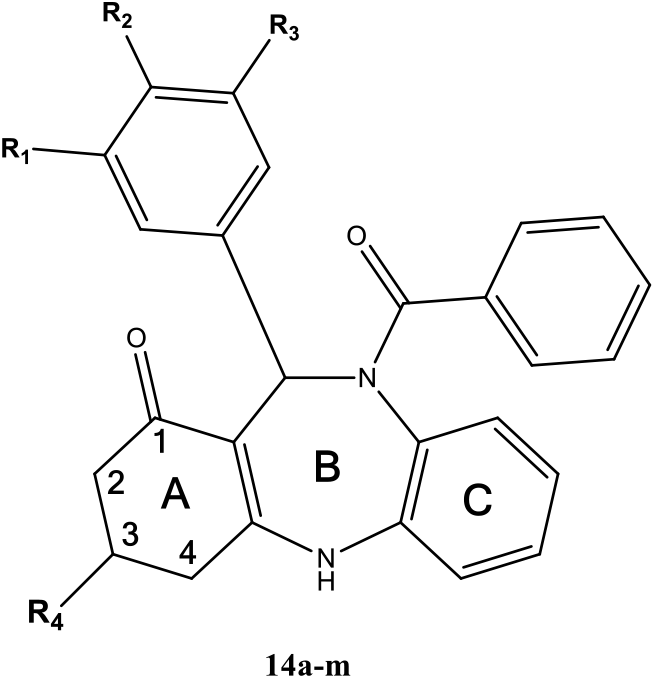

**Tab. 6.**
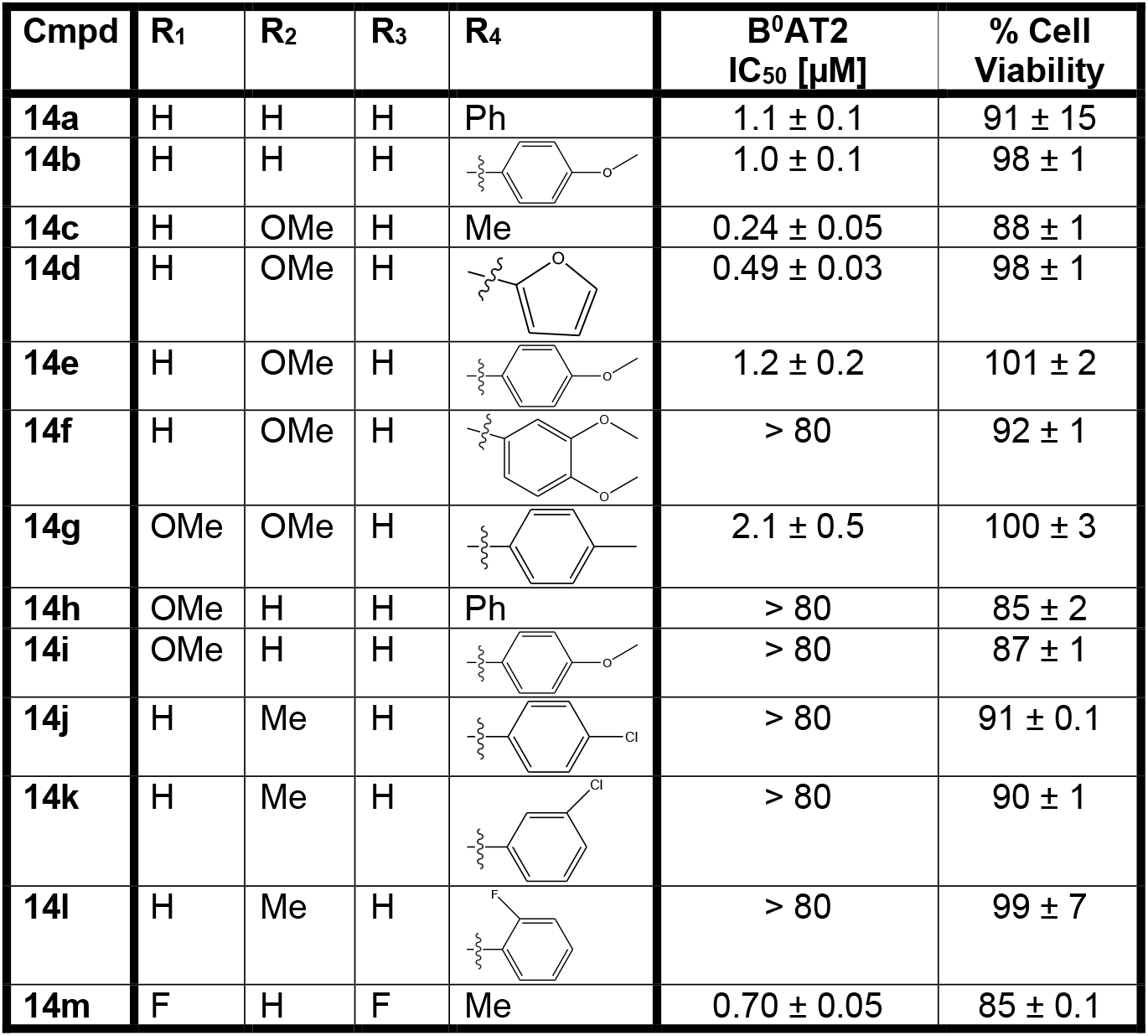

| Cmpd | R<sub>1</sub> | R<sub>2</sub> | R<sub>3</sub> | R<sub>4</sub> | B<sup>0</sup>AT2<br>IC<sub>50</sub> [μM] | % Cell<br>Viability |
| --- | --- | --- | --- | --- | --- | --- |
| **14a** | H | H | H | Ph | 1.1 ± 0.1 | 91 ± 15 |
| **14b** | H | H | H |  | 1.0 ± 0.1 | 98 ± 1 |
| **14c** | H | OMe | H | Me | 0.24 ± 0.05 | 88 ± 1 |
| **14d** | H | OMe | H |  | 0.49 ± 0.03 | 98 ± 1 |
| **14e** | H | OMe | H |  | 1.2 ± 0.2 | 101 ± 2 |
| **14f** | H | OMe | H |  | > 80 | 92 ± 1 |
| **14g** | OMe | OMe | H |  | 2.1 ± 0.5 | 100 ± 3 |
| **14h** | OMe | H | H | Ph | > 80 | 85 ± 2 |
| **14i** | OMe | H | H |  | > 80 | 87 ± 1 |
| **14j** | H | Me | H |  | > 80 | 91 ± 0.1 |
| **14k** | H | Me | H |  | > 80 | 90 ± 1 |
| **14l** | H | Me | H |  | > 80 | 99 ± 7 |
| **14m** | F | H | F | Me | 0.70 ± 0.05 | 85 ± 0.1 |

### 4.4. Characterization of representative 1,5-benzodiazepine analogs

A preliminary ADME profiling of the most potent 1,5-benzodiazepine analogs showed that compound **14c** had promising physicochemical properties (Fig. 3a) and was thus characterized in more detail.

**Fig. 3:**
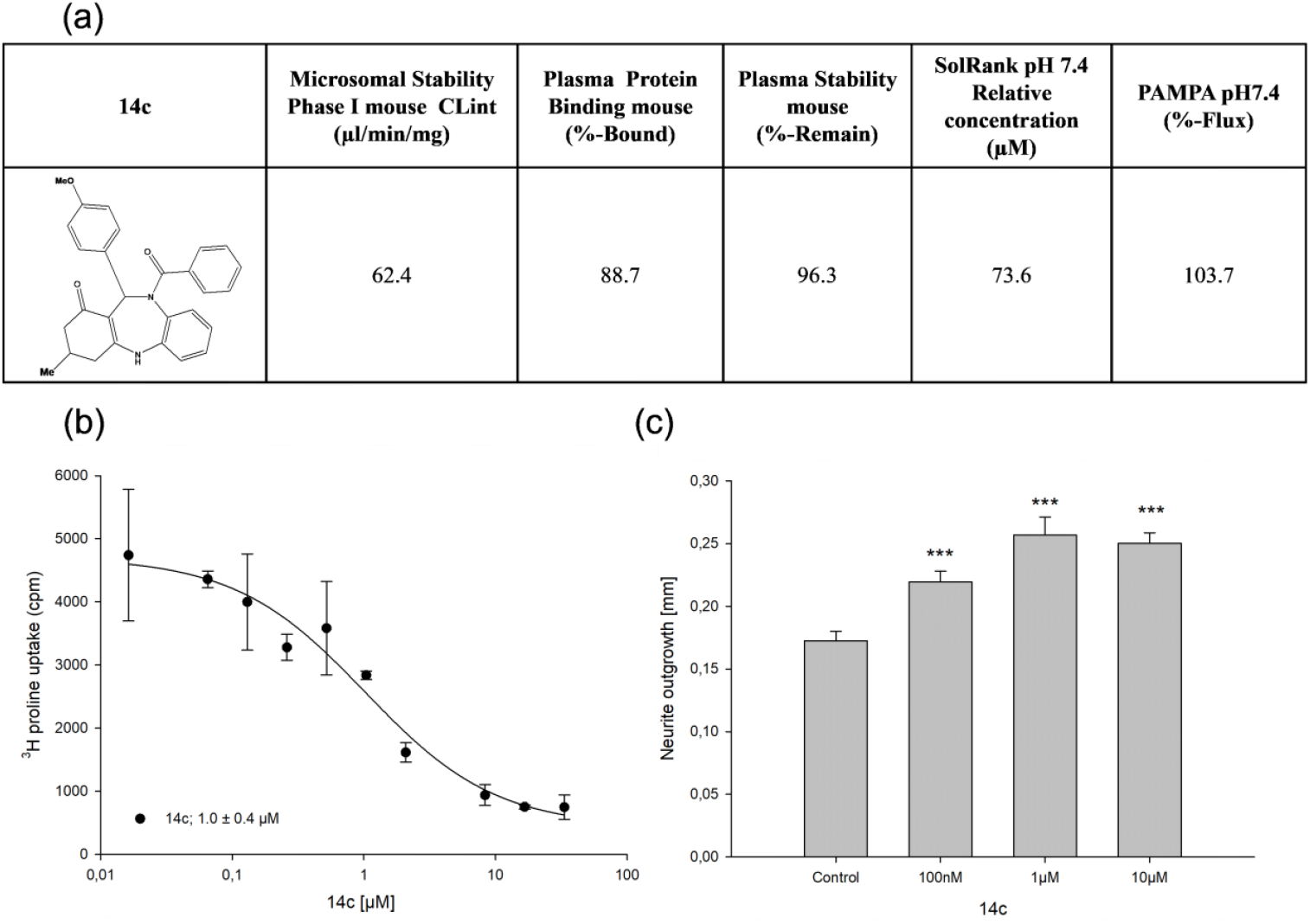
Characterization of the 1,5-benzodiazepine **14c**. (a) Structure and in vitro ADME parameters of **14c**. (b) **14c** dose-dependently inhibited ^3^H-proline uptake in cultured primary hippocampal neurons overexpressing B^0^AT2. (c) **14c** dose-dependently enhanced neurite outgrowth in cultured primary hippocampal.

To confirm the activity of the 1,5-benzodiazepine analogs for untagged B^0^AT2 in a relevant environment, selected representatives were tested in cultured primary neurons that overexpressed B^0^AT2/SLC6A15. Compounds **11x, 14c** and **14d** inhibited ^3^H-proline uptake in a dose-dependent manner (Fig. 3b and Fig. S1a). The potency was comparable to the values obtained in eGFP-hSLC6A15-overexpressing HEK293 cells confirming the latter as a viable model for the identification of B^0^AT2 inhibitors.

Finally, we evaluated the effect of prolonged treatment of 1,5-benzodiazepine analogs on neuronal integrity. As a functional readout, we used neurite outgrowth as a cellular model for neuronal differentiation. Surprisingly, compounds **11x, 14c** and **14d** all increased neurite length of dissociated hippocampal neurons in a dose-dependent manner (Fig. 3b, Fig. S1b and S1c), pointing to an hitherto unappreciated role of B^0^AT2 in neuronal dynamics.

#### Discussion and conclusions

The SLC6A family is a highly privileged target class for psychoactive drugs. This can be attributed to the pivotal role that many of the SLC6A transporters play in regulation neuronal transmission and to their high druggability, i.e., the comparatively high likelihood to identify drug-like ligands for them. In our screening we could readily identify several different classes of B^0^AT2 inhibitors. This demonstrates that like other SLC6A transporters B^0^AT2 is a very druggable target.

So far the best available B^0^AT2 inhibitor has been pipeolic acid, a competitive substrate with an IC_50_ of 900 µM (Broer *et al*., 2006). Our best inhibitors like compound **14c** are > 1000-fold more potent, have promising physicochemical properties and an excellent selectivity profile, allowing studies in advanced cellular systems and possibly even animal models.

A rather surprising aspect of the identified B^0^AT2 inhibitors is their extraordinary selectivity within the SLC6A family. None of the identified B^0^AT2 inhibitors inhibited SERT and only compound **7** displayed 100-fold weaker inhibition of GAT and DAT. Most surprising, however, is the exquisite selectivity over NTT4/SLC6A17, the closest homolog of B^0^AT2. NTT4 and B^0^AT2 have a very similar expression pattern in the brain and they transport the same amino acid substrates (Parra *et al*., 2008; Zaia *et al*., 2009). These two neutral amino acid transporters share 66 % overall sequence identity but are completely identical within 8Å of the putative primary substrate site (Bröer *et al*., 2012; Yamashita *et al*., 2005; Zaia *et al*., 2009). Of the identified B^0^AT2 inhibitors only two compounds (**2a** and **7**) showed cross-reactivity at NTT4.

The remarkable subtype selectivity raises questions about the binding site of the identified B^0^AT2 inhibitors. The cocrystal structures of the bacterial amino acid transporter LeuT, a biochemically model system for mammalian SLC6A transporters, revealed two potential binding sites, one located at the substrate binding site and another in an allosteric pocket above the latter (Cuboni *et al*., 2014; Penmatsa *et al*., 2013; Singh *et al*., 2007; Yamashita *et al*., 2005; Zhou *et al*., 2007). A detailed kinetic analysis of the identified inhibitors and mutagenesis studies, e.g. of the two potential binding sites, are required to shed light on the binding mode of the identified B^0^AT2 inhibitors.

Compounds corresponding to the motif defined by the scaffolds **10-14** have not been described in the literature. The closest compounds are 1,5-benzodiazepines that have been developed as inhibitors of the hepatitis C virus 5B RNA-dependent RNA polymerase (McGowan *et al*., 2009; Nyanguile *et al*., 2008; Vandyck *et al*., 2009). This suggests that this type of compounds can be optimized into advanced drug-like compounds.

B^0^AT2 is expressed predominantly in the brain but so far its cellular role in neurons is poorly understood. Among other hypotheses, the removal of potentially neuroactive amino acids like proline from the synaptic cleft or the supply of branched amino acids as precursors for the synthesis of neurotransmitters have been discussed (Broer, 2013). Our serendipitous discovery that several B^0^AT2 inhibitors enhance neurite outgrowth in neurons suggests that B^0^AT2 might play an unanticipated in neuronal differentiation and in neuronal remodelling. The mechanistic details and the physiological relevance of this cellular function are currently being explored.

## Abbreviations

GABA: *γ-*aminobutyric acid
SERT: serotonin transporter
NET: norepinephrine transporter
DAT: dopamine transporter
GAT1: GABA transporter 1
GlyT1: glycine transporter 1

## Acknowledgements

We are indebted to Drs. Blakely, Wanner and Reimer for plasmids or cell lines for SERT, GAT1 or NTT4-B^0^AT2. We thank C. Sippel and Dr. A. Kirschner for technical support, Drs S. Santarelli and M. Schmidt for providing primary hippocampal embryonic neurons, Drs. A. Wolf, M. Baumann, T. Bergbrede, and S. Sievers for support and access to the high-throughtput facilities and the COMAS library, and Dr. F. Holsboer for institutional support. This work was funded in part by the HolsboerMaschmeyer NeuroChemie GmbH.

---

**Fig. S1:**
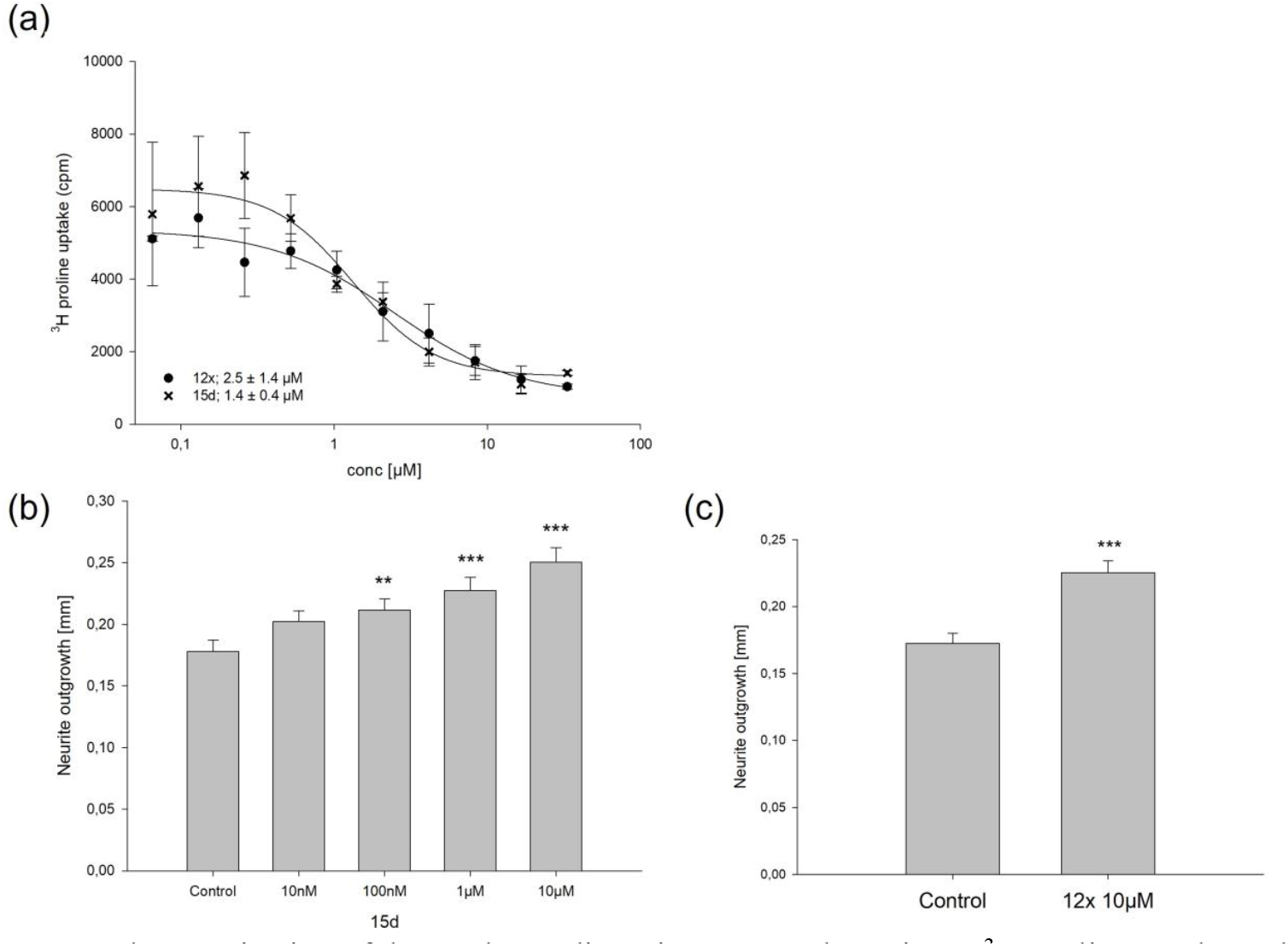
Characterization of the 1,5-benzodiazepines **11x** and **14d** in (a) ^3^H-proline uptake and (b, c) neurite outgrowth in cultured primary hippocampal neurons.

